# A single locus determines praziquantel response in *Schistosoma mansoni*

**DOI:** 10.1101/2023.11.01.565202

**Authors:** Frédéric D. Chevalier, Winka Le Clec’h, Matthew Berriman, Timothy J.C. Anderson

## Abstract

We previously performed a genome-wide association study (GWAS) to identify the genetic basis of praziquantel (PZQ) response in schistosomes, identifying two quantitative trait loci (QTL) situated on chromosome 2 and chromosome 3. We reanalyzed this GWAS using the latest (v10) genome assembly showing that a single locus on chromosome 3, rather than two independent loci, determines drug response. These results reveal that praziquantel response is monogenic and demonstrates the importance of high-quality genomic information.

## TEXT

In a recent research article (1), we performed a genome-wide association study (GWAS) to identify the genetic basis of praziquantel (PZQ) response in schistosomes. This study leveraged a mixed population of PZQ-resistant and PZQ-sensitive *Schistosoma mansoni* generated by laboratory selection (SmLE-PZQ-R) (2), and we compared genome-wide allele frequencies in populations of PZQ-treated and untreated adult worms to determine the genetic basis of PZQ response. We identified two quantitative trait loci (QTL) associated with drug response situated on chromosome 2 and chromosome 3 (Fig. 1A). On chromosome 3, we determined that the *Sm*.*TRPM*_*PZQ*_ gene, which encodes a transient potential receptor channel, was the cause of variation in PZQ response. This conclusion was further supported by an independent study using pharmacological approaches (3). In addition, we showed lower *Sm*.*TRPM*_*PZQ*_ gene and isoform expression in PZQ-resistant (PZQ-R) worms, suggesting that expression level may determine the PZQ response phenotype. However, we were unable to identify a causative gene within the chromosome 2 QTL. In this note, we revisit the published dataset using an updated and improved genome assembly to further investigate the chromosome 2 QTL, to try to resolve the inconsistencies observed in Le Clec’h *et al*. (1), and to better understand the genetic architecture of PZQ response.

**Fig. 1.**
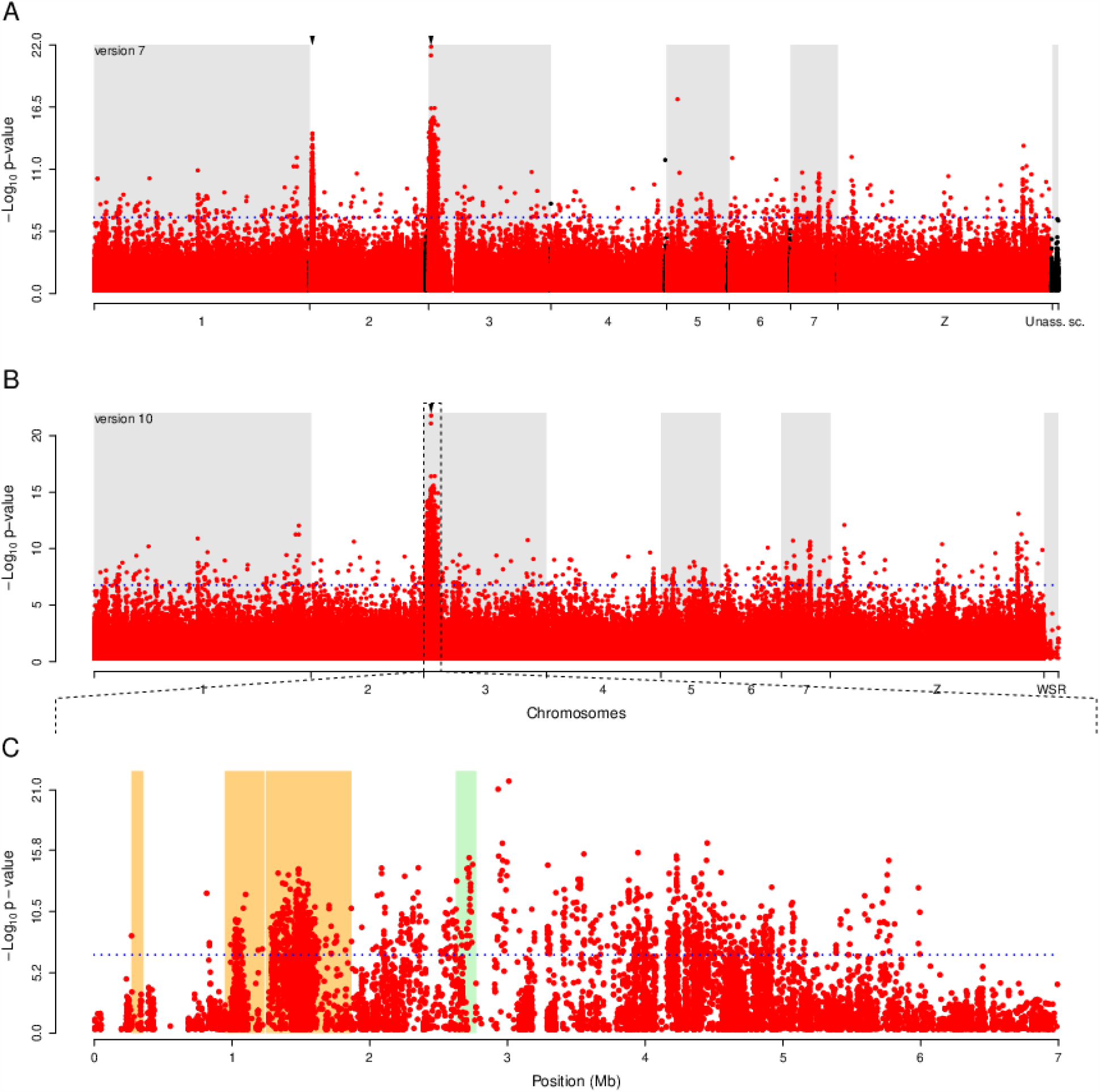
Comparison of the genome-wide association mapping of PZQ response against versions 7 and 10 of the *S. mansoni* reference genome. The Manhattan plot identifies genome regions that differ in allele frequency between PZQ sensitive and resistant worm pools. While the mapping done with the version 7 of the reference genome revealed two QTL peaks on chromosomes 2 and 3 **(A)**, mapping against the version 10 of the genome highlighted a single peak on chromosome 3 **(B)**. The QTL on chromosome 2 was actually an artifact due to misplaced sections of the chromosome 3 (orange box) **(C)**. Blue dotted line refers to the Bonferroni significance threshold; red and black dots represent association of individual SNPs for assembled and unassembled scaffolds respectively; green box marks the position of the Sm.TPRM_PZQ_ gene.

We used the version 7 of the *S. mansoni* genome to map our genomic data in Le Clec’h *et al*. (1). This version contained 312 unassembled scaffolds, many of which represented alternative haplotypes for regions across the genome, in addition to the near complete assembly of the 7 autosomes and ZW sex chromosomes. However, the most recent of the *S. mansoni* reference genome assemblies (version 9 (4) and version 10) have been substantially improved. The latest version 10 of the genome (https://parasite.wormbase.org/Schistosoma_mansoni_prjea36577; (5)) is now assembled in full-length chromosomes. The unassembled scaffolds have been either incorporated into the main chromosomal assemblies, or removed as haplotypic alternatives for many regions. Importantly some chromosomal regions have been reassigned to their correct chromosomal locations (either on the same chromosome or on a different chromosome). These assembly changes have particularly affected the contiguity of the sex chromosomes but have also affected regions of chromosome 2. In parallel with the assembly revision, gene models have been further improved in version 10 to better reflect experimental evidence; predicted transcript isoforms are now only included when alternative intron:exon boundaries are supported by ≥10% alignment depth from a single RNA sequencing library. This gene model curation has resulted in a global reduction in transcript isoforms predicted: while 27.58% of gene models encoded more than one isoform in version 7, this number fell to 8.65% in version 10. The *Sm*.*TRPM*_*PZQ*_ is one of the genes affected, with a reduction in number of major annotated isoforms from 7 to 3. With the release of this revised and improved reference genome, we took the opportunity to revisit our genetic mapping experiments to re-evaluate the genetic architecture of PZQ response in schistosomes. We also reanalyzed our transcriptomic data to determine whether *Sm*.*TRPM*_*PZQ*_ gene expression remains associated with differences in PZQ response.

Our revised genetic mapping using the latest version of the *S. mansoni* reference genome revealed a single QTL on chromosome 3 associated with PZQ response. The chromosome 2 QTL is no longer present (Fig. 1B). This change is a direct consequence of the relocation of sections of chromosome 2 to chromosome 3 (Fig. 1C). This is reflected by the size of the new QTL (5,723,424 bp), which is approximately the sum of the two previous QTLs (chr. 2: 1,166,271 bp; chr. 3: 3,990,733 bp; total: 5,157,004 bp). The chromosome 2 QTL is now located, in three segments, on the boundary of the chromosome 3 QTL, near the beginning of the p-arm of the chromosome. The location on the boundary of the chromosome 3 QTL is likely to explain the absence of correlation between phenotype and genotype previously shown using individual worms (1). The relocation of chromosome 2 QTL increased the total number of genes under the updated chromosome 3 QTL from 91 to 137, of which 125 are expressed in adults. The strongest associated SNP marker (position 3,010,821 T > C) was still located in the *SOX13* transcription factor gene (Smp_345310). In addition, the two deletions previously identified close to the *Sm*.*TRPM*_*PZQ*_ and the *SOX13* genes and associated with the resistant phenotype were confirmed by remapping to the version 10 genome (positions 2,775,001 - 2,900,000 and 3,175,001 - 3,300,000, respectively).

We revisited our transcriptomic analysis using the updated annotation produced alongside the version 10 of the reference genome. This updated annotation included a revised *Sm*.*TRPM*_*PZQ*_ gene model with minor changes in number of exons but extensive alterations in number of isoforms (Fig. 2A). The total number of exons was reduced from 41 to 38, with modifications to the boundaries of some exons. As expected, these minor changes had a limited impact on the overall gene expression. Our revised transcriptomic analysis confirmed the reduced expression of *Sm*.*TRPM*_*PZQ*_ gene in the SmLE-PZQ-ER parasites (enriched for PZQ-R allele) (Fig. 2B-C) in adult and juvenile worms of both sexes. We confirmed that the *Sm*.*TRPM*_*PZQ*_ gene is the only gene under the QTL with a significant change in expression (Fig. 2B). The gene has a lower expression in SmLE-PZQ-ER adult male and female worms compared to their SmLE-PZQ-ES counterparts (Fig 2C). We also confirmed very low expression in adult females of both SmLE-PZQ-ER and -ES and the higher gene expression pattern in SmLE-PZQ-ES juvenile worms.

**Fig. 2.**
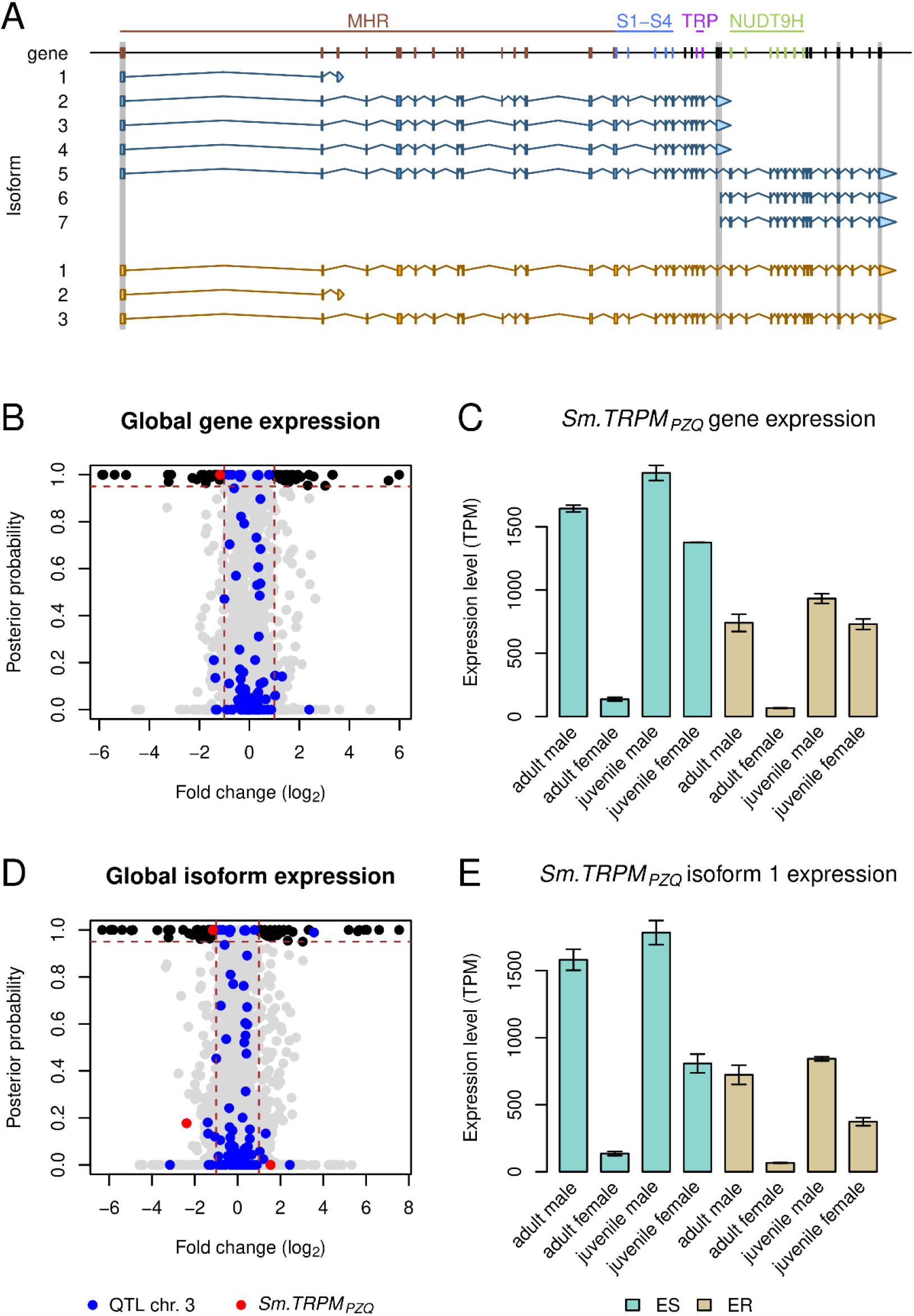
Gene expression differences between SmLE-PZQ-ES and SmLE-PZQ-ER parasites. **(A)** The gene and isoform annotations were revised with the version 10 of the reference genome, which led to an updated *Sm*.*TRPM*_*PZQ*_ gene model. The number of predicted isoforms changed from 7 to 3: isoform 3, 4, 6, and 7 were discarded; isoforms 2 (now 3) and 5 (now 1) were updated. Changes in exons (either removal or updates of their boundaries) are highlighted with grey boxes. Exons encoding major protein domains are highlighted on the gene model: the TRPM homology region (MHR) domain (brown), the four transmembrane-spanning helices (S1–S4) (blue) and TRP box (purple), which are key in the interaction between the channel and PZQ, and the NUDT9H domain (green). **(B)** The *Sm*.*TRPM*_*PZQ*_ gene is the only gene under the chromosome 3 QTL to show differential expression between adult male PZQ-ES and PZQ-ER. **(C)** *Sm*.*TRPM*_*PZQ*_ gene showed reduced expression in PZQ-ER parasites at all stages and for both sexes. Gene expression is overall higher in juveniles than adults, while gene expression in adult females is very low, likely explaining their natural resistance to PZQ. **(D)** Among the three isoforms of the *Sm*.*TRPM*_*PZQ*_ gene, only isoform 1 showed differential expression. **(E)** The expression pattern of this isoform is very similar to the gene expression pattern, with high expression in naturally resistant juveniles. On the volcano plots, black dots shows genes or isoforms with significant differential expression genome-wide, blue dots shows genes or isoforms located under the chromosome 3 QTL, red dots shows *Sm*.*TRPM*_*PZQ*_ gene or isoform, vertical brown lines show a 2-fold threshold in differential expression, horizontal brown line shows a threshold of 0.95 posterior probably in differential expression.

However, the updated analysis of isoform expression differed in several ways from that previously observed. The revised gene models resulted in reduction of annotated isoforms from 7 (version 7) to 3 (version 10) (Fig. 2A). Specifically, isoforms 3, 4, 6 and 7 from version 7 are no longer present. Isoform 1 is the most abundant of the 3 isoforms, accounting for the majority (88.32-100%) of *Sm*.*TRPM*_*PZQ*_ trancripts in adults. We detected reduced expression of isoform 1 (formerly isoform 5) in SmLE-PZQ-ER adult males compared to SmLE-PZQ-ES adult males. SmLE-PZQ-ER adult males show comparable expression of isoform 1 to SmLE-PZQ-ES juvenile females which are naturally resistant. The new annotation of the version 10 genome does not exclude the possibility of rarer exon combinations, but the simplified isoform profile, and the predominance of isoform 1, now allows us to focus on this isoform for future analyses. We previously hypothesized that isoform 6 expression might be associated with PZQ sensitivity because of its high expression only in SmLE-PZQ-ES adult males (1). Isoform 6 corresponded to the terminal 15 exons of the gene model but, from existing short-read data, there is insufficient evidence to infer an appropriate transcriptional start for this truncated isoform. The isoform, therefore, no longer exists in the version 10 assembly and the association is likely to be spurious and allowing us to reject this hypothesis.

Our reanalysis now clearly shows that PZQ response is a single gene recessive trait in the laboratory populations studied and does not involve two independent loci as previously indicated (1). This is consistent with the observation that drug resistance typically has a simple genetic basis and is often monogenic (6). If PZQ resistance is also monogenic in natural parasite populations this will greatly simplify molecular monitoring in control programs. Reanalysis of the transcript data using version 10 annotation also simplifies our understanding of this system, showing that just isoform 1 (of 3 isoforms) predominates, and shows differential expression between resistant and sensitive parasites. Future work can focus on how expression of isoform 1 impacts drug response. These updated results, utilizing the latest high-quality *S. mansoni* reference genome, underscore the importance of a robust reference genome for precise genetic mapping of critical biomedical traits, such as drug resistance in pathogens. Similar efforts to enhance genome assemblies have also been recently undertaken for the two other major schistosome species, *S. haematobium* and *S. japonicum* (7, 8). High-quality genomes have also improved our understanding of drug resistance in *Haemonchus contortus*, a major gastrointestinal nematode of small ruminants (9). Our present results serve as a compelling illustration of the ongoing need for continuous improvements in genome assembly, after the initial publication (10).

## Supplementary Materials

Table S1. Genes under QTL of chromosome 3.

## Acknowledgments

This research was supported by a Cowles fellowship (WLC) from Texas Biomedical Research Institute (13-1328.021), and NIH R01AI133749 (TJCA) and was conducted in facilities constructed with support from Research Facilities Improvement Program grant C06 RR013556 from the National Center for Research Resources. SNPRC research at Texas Biomedical Research Institute is supported by grant P51 OD011133 from the 570 Office of Research Infrastructure Programs, NIH. The authors thank Sarah Buddenborg and Steve Doyle for early access to unpublished sequence and revised annotation, as well as Duncan Berger, for the original observation of a misplaced section of chromosome 3.

## Funding

National Institutes of Health grant <u>1R01AI123434</u> (TJCA) National Institutes of Health grant 1R01AI133749 (TJCA)

## Author contributions

Conceptualization: FDC, WLC, TJCA

Methodology: FDC, WLC, TJCA, MB

Investigation: FDC

Visualization: FDC

Funding acquisition: TJCA

Project administration: TJCA

Supervision: TJCA, WLC, FDC

Writing – original draft: FDC

Writing – review & editing: TJCA, WLC, FDC, MB

## Competing interests

Authors declare that they have no competing interests.

## Data and materials availability

Sequence data is available from: PRJNA699326, PRJNA701978, PRJNA704646

Code is available on Github (https://github.com/fdchevalier/PZQ-R_DNA-seq, https://github.com/fdchevalier/PZQ-R_RNA-seq)

